# Impact of viral membrane oxidation on SARS-CoV-2 spike protein transmembrane anchoring stability

**DOI:** 10.64898/2026.03.27.714475

**Authors:** Maryam Ghasemitarei, Cristina Gyursánszky, Mikko Karttunen, Tapio Ala-Nissila

**Affiliations:** MSP Group, Department of Applied Physics, Aalto University, P.O. Box 15600, FI-00076 Aalto, Espoo, Finland; Department of Technical Physics, University of Eastern Finland, P.O. Box 1627, FI-70211 Kuopio, Finland; European Laboratory for Learning and Intelligent Systems (ELLIS) Institute Finland, Maarintie 8, 02150 Espoo, Finland; Department of Mathematical Sciences and Interdisciplinary Centre for Mathematical Modelling, Loughborough University, Loughborough LE11 3TU, United Kingdom

**Keywords:** SARS-CoV-2 spike protein, Lipid peroxidation, POPC oxidation, Molecular dynamics simulation, Potential of mean force, Membrane stability

## Abstract

Reactive oxygen species generated during inflammation can oxidize viral envelope lipids, with outcomes ranging from modulated infectivity to viral inactivation. For SARS-CoV-2, the molecular mechanisms by which membrane lipid oxidation influences spike protein anchoring remain poorly understood. We use all-atom molecular dynamics (MD) simulations to quantify how graded oxidation of 1-palmitoyl-2-oleoyl-*sn*-glycero-3-phosphocholine (POPC) affects the anchoring of the SARS-CoV-2 spike transmembrane (TM) region in an endoplasmic-reticulum–Golgi intermediate compartment (ERGIC)-like multicomponent membrane. Viral envelopes containing 0, 25, 50, 75, and 100% oxidized POPC (PoxnoPC) corresponding to 0 − 55% oxidation of all PO-type phospholipids were simulated with the spike TM helix and cytoplasmic tail embedded in a POPC/POPE/POPI/POPS/cholesterol mixture. Steered MD and umbrella sampling were used to calculate the potential of mean force (PMF) for extracting the TM+CT region along the membrane normal. Partial oxidation (25 − 75% POPC) produced reductions in the detachment barrier that were not statistically distinguishable from the native system within the sampling uncertainty, whereas full POPC oxidation lowered the anchoring free energy by about 23% (from 606 ± 39 to 464 ± 38 kJ mol^−1^), indicating that oxidation of roughly half of the glycerophospholipids can measurably weaken spike-membrane coupling. Despite this reduction, the remaining barrier (about 180*k*_B_*T* ) is still large, suggesting that oxidation alone may be insufficient for spontaneous spike detachment and likely acts synergistically with mechanical forces during fusion or immune engagement. Analysis of acyl-chain order parameters, area per lipid, membrane thickness, number-density profiles, and lateral lipid clustering reveals that POPC peroxidation decreases lipid order, thins and softens the bilayer, and disrupts cholesterol-stabilized clusters that refer to large cooperative lipid assemblies (>10 lipids) identified via RDF-based clustering. These oxidation-induced changes reduce hydrophobic matching around the TM helix and facilitate its extraction from the viral envelope. Our results provide a mechanistic link between lipid peroxidation, membrane nanostructure, and spike anchoring, supporting lipid oxidation for example during cold atmospheric plasma or ozone treatment as a physically grounded contributing antiviral mechanism against SARS-CoV-2.

## 1. Introduction

Reactive oxygen species (ROS) generated during immune responses can induce oxidative damage to biomolecules, including lipids, proteins, and nucleic acids. Although oxidative stress is classically viewed as a host defense mechanism, many viruses have evolved strategies to tolerate or even exploit oxidizing environments for their replication and propagation [1, 2]. SARS-CoV-2 infection, for instance, is known to activate NADPH oxidase in endothelial and epithelial cells, increasing reactive oxygen species (ROS) production and inducing mitochondrial dysfunction. This oxidative stress drives endothelial dysfunction, contributing to COVID-19 pathogenesis and cardiovascular complications including thrombosis and atherosclerosis [3, 4]. Among the macromolecules susceptible to ROS-mediated damage, polyunsaturated fatty acids in membrane lipids are particularly prone to oxidation. Peroxidation of unsaturated fatty acyl chains generates lipid hydroperoxides and reactive aldehydes [5, 6] which can profoundly alter membrane structure and biophysical properties. Oxidized lipids disorder membrane structure and enhance permeability, rendering the bilayer more susceptible to mechanical stress-induced pore formation [7–9], thereby modifying the functions of embedded proteins, both by altering the biophysical properties of the lipid bilayer and through direct covalent modification by reactive aldehyde byproducts. In cellular contexts, low levels of lipid peroxidation can modulate signaling pathways and membrane properties, whereas high levels cause loss of membrane integrity and cell death [10].

Lipid oxidation is therefore implicated in diverse pathological conditions, including atherosclerosis, diabetes, cancer, infectious diseases, and neurodegenerative disorders [5, 11], but has also been exploited therapeutically, for example in cold atmospheric plasma (CAP) treatments in oncology and antiviral applications [6, 7, 12]. For enveloped viruses such as SARS-CoV-2, which derive their membrane from the host, oxidative modification of viral lipids can have dual and dosedependent effects. Mild oxidative environments may alter membrane fluidity or protein conformational dynamics in ways that subtly enhance viral fusion. In contrast, extensive lipid peroxidation compromises envelope integrity and inactivates the virus. Recent studies support this dichotomy: moderate oxidative modifications can strengthen SARS-CoV-2 receptor binding and increase viral entry [6, 13], whereas severe oxidation such as that during CAP exposure or ozone treatment fragments the spike protein, disrupts the viral membrane, and abolishes infectivity [14–16].

The spike (S) glycoprotein, the primary determinant of SARS-CoV-2 host-cell entry, is highly sensitive to its membrane environment. Spike exists as a trimer embedded in the viral envelope and undergoes transitions between “down” and “up” conformations that regulate accessibility of the receptor-binding domain (RBD) for ACE2 engagement [17]. Modulation of membrane composition including lipid unsaturation, cholesterol content, anionic lipids, or covalent modifications such as palmitoylation can significantly influence spike stability, orientation, flexibility, and fusion competence. Oxidative changes in membrane lipids therefore represent a plausible but underexplored mechanism for regulating spike conformational stability and its interactions with the host membrane during the fusion process.

Despite growing evidence linking oxidative stress, lipid peroxidation, and SARS-CoV-2 biology, the molecular-level effect of oxidized membrane lipids on spike stability remains largely unknown. Experimental studies indicate that lipid peroxides can modulate viral fusion efficiency, but the atomistic details on how oxidized membrane lipids alter bilayer order, influence spike anchoring, or destabilize the transmembrane region are not yet established. SARS-CoV-2 assembly and budding occur in the endoplasmic reticulum–Golgi intermediate compartment (ERGIC), where the viral envelope is obtained from host intracellular membranes [18, 19]. These intracellular membranes are rich in phosphatidylcholine (PC), one of the dominant phospholipid classes in mammalian cells [20]. Among PC molecular species, POPC (1-palmitoyl-2-oleoyl-sn-glycero-3-phosphocholine) is one of the most prevalent in eukaryotic membranes and is widely used as a representative zwitterionic lipid in biophysical studies [21, 22]. In some SARS-CoV-2 membrane models, POPC constitutes a significant fraction of the viral envelope lipids [23]. POPC contains a monounsaturated oleoyl chain, making it highly susceptible to ROS-induced peroxidation [5, 8, 11].

Oxidized POPC derivatives such as PoxnoPC are well-characterized lipid peroxidation products and are extensively used to model oxidative membrane damage [24]. PoxnoPC forms via cyclization and cleavage of a POPC peroxyl radical intermediate, yielding two aldehyde-containing fragments, of which PoxnoPC is the dominant product [7]. The aldehyde form is both more stable and more experimentally abundant than the corresponding hydroperoxide [6]. Oxidation of POPC leads to pronounced changes in membrane biophysics including bilayer thinning and enhanced permeability, while effects on acyl-chain order and area per lipid are more complex and depend on the specific oxidation product [8, 24]. In a recent work, Xie et al. [25] have provided mechanistic insight into how truncated oxidized phospholipids, including PoxnoPC, disrupt membrane structure. They demonstrated that the membranolytic activity depends on both specific chemical structure, with aldehydes and carboxylic acids exhibiting distinct permeabilization efficiencies, and the pH of the environment. Notably, ionization of the terminal carboxyl group at physiological pH increases the intrinsic molecular curvature of the oxidized lipid, promoting membrane curvature and permeabilization, whereas the aldehyde form of PoxnoPC does not induce such curvature but instead enhances permeability to larger molecules. These findings underscore that the biophysical consequences of lipid oxidation are not uniform but are instead governed by the precise molecular identity of the oxidation products and the local chemical environment. Because POPC is biologically abundant and computationally well-parameterized, focusing on POPC oxidation provides a robust and interpretable framework for evaluating how lipid peroxidation may alter spike–membrane stability in SARS-CoV-2. However, we emphasize that biological lipid-peroxidation generates a complex mixture of products including hydroperoxides, epoxides, hydroxides, and carboxylic acid-terminated fragments, each with distinct effects on membrane structure [25]. By modeling only the aldehyde product PoxnoPC, this study captures one specific axis of the peroxidation landscape, and the generalizability of our conclusions to other oxidation products remains to be established.

The decision to oxidize only POPC, rather than all PO-type glycerophospholipids included in the SARS-CoV-2 membrane model (POPE, POPI, and POPS), is motivated by both compositional and biophysical considerations. All PO-type lipids share the same acyl-chain architecture—a saturated palmitoyl (16:0) chain at the *sn*-1 position and a monounsaturated oleoyl (18:1) chain at the *sn*-2 position—so their differences arise primarily from their headgroups. POPC provides a natural axis for defining systematic oxidation levels while maintaining overall membrane composition consistent with previous SARS-CoV-2 models [23]. Furthermore, the zwitterionic, cylindrical PC headgroup forms fluid bilayers with minimal intrinsic curvature, allowing oxidation-induced changes to be attributed primarily to acyl-chain chemistry rather than headgroup effects [26]. In contrast, POPS and POPI are anionic minor components that participate in specific electrostatic interactions and possess non-cylindrical headgroups whose oxidation would simultaneously alter membrane charge and curvature [27, 28]. POPE, with its small ethanolamine headgroup, forms strong inter-lipid hydrogen bonding and promotes negative curvature [29], making its oxidation more difficult to isolate from headgroup-driven structural effects. By oxidizing only POPC whose acyl chain structure matches that of the other PO lipids while having a neutral, well-characterized headgroup, we specifically probe the impact of chain oxidation without introducing confounding variations in membrane charge or curvature. Additionally, POPC/PoxnoPC is among the most extensively parameterized oxidized lipid systems in atomistic MD, enabling direct comparison with prior computational and experimental studies.

Since lipid oxidation increases membrane fluidity, permeability, and reduces stiffness, we hypothesize that it may also weaken the anchoring and stability of the SARS-CoV-2 spike protein within the viral membrane. Such destabilization could indirectly influence the spike’s ability to engage ACE2, a necessary step for mediating membrane fusion and viral entry [17]. Consistent with this hypothesis, oxidation of viral membranes has been shown to inhibit late-stage fusion in other enveloped viruses. For example, Vigant et al. demonstrated that membrane-embedded photosensitizers generate singlet oxygen and selectively oxidize viral lipids, leading to impaired fusion in HIV and Nipah virus [12]. These findings highlight that oxidative modification of viral envelopes can act as an antiviral mechanism. However, the outcome of lipid oxidation strongly depends on membrane composition, particularly on cholesterol content and lipid diversity [5, 7]. Given that the SARS-CoV-2 envelope contains moderate levels of cholesterol and a multicomponent lipid mixture, it remains unclear how increasing POPC oxidation will affect spike–membrane stability. This motivates the present work.

In the present study, we investigate how increasing levels of POPC oxidation influence the stability of the SARS-CoV-2 spike protein within the viral membrane. We construct membrane systems containing 0%, 25%, 50%, 75%, and 100% oxidized POPC, corresponding to 0 − 55% oxidation of the total PO-type phospholipids (POPC, POPE, POPI, POPS), and perform extensive all-atom molecular dynamics (MD) simulations of the transmembrane region of the spike protein embedded in each membrane. To quantify spike–membrane anchoring strength, we carry out steered molecular dynamics (SMD) pulling of the spike transmembrane region, followed by umbrella sampling (US) [30] and calculation of the potential of mean force (PMF) along the extraction pathway. In addition to free-energy analysis, we characterize how lipid peroxidation alters membrane biophysical properties. Together, these simulations provide a molecular-level framework for understanding how lipid oxidation modifies membrane structure and modulates spike anchoring in SARS-CoV-2.

## 2. Material and method

### 2.1. System setup

The SARS-CoV-2 spike protein is a large class I viral fusion protein composed of 1273 amino acids per monomer. However, available cryo-EM structures resolve only the ectodomain (residues 1 − 1146) and do not include the transmembrane helix (TM), the membrane-embedded region, or cytoplasmic tail (CT). To model the membrane-embedded portion of the spike, we used the fulllength, computationally reconstructed spike model developed by Woo et al. [31], which provides atomistic coordinates for the HR2 region, TM helix, and CT domain. A schematic representation of the protein–membrane systems is shown in Figure 1. We note that an experimental NMR structure of the SARS-CoV-2 spike TM domain trimer in lipid bicelles has, since the work of Woo et al. [31], become available by Fu and Chou [32]. However, in that study the structure was determined for a short peptide carrying four explicit point mutations in DMPC/DH_6_PC bicelles. The Woo et al. model was chosen here because it provides wild-type sequence coordinates for the complete continuous membrane-proximal stalk, transmembrane, and cytoplasmic regions (residues 1168–1273) needed to capture the necessary thermodynamic anchors for PMF calculations. The sensitivity of the computed PMF to the precise TM trimer packing geometry remains to be established and is beyond the scope of the current study.

**Figure 1:**
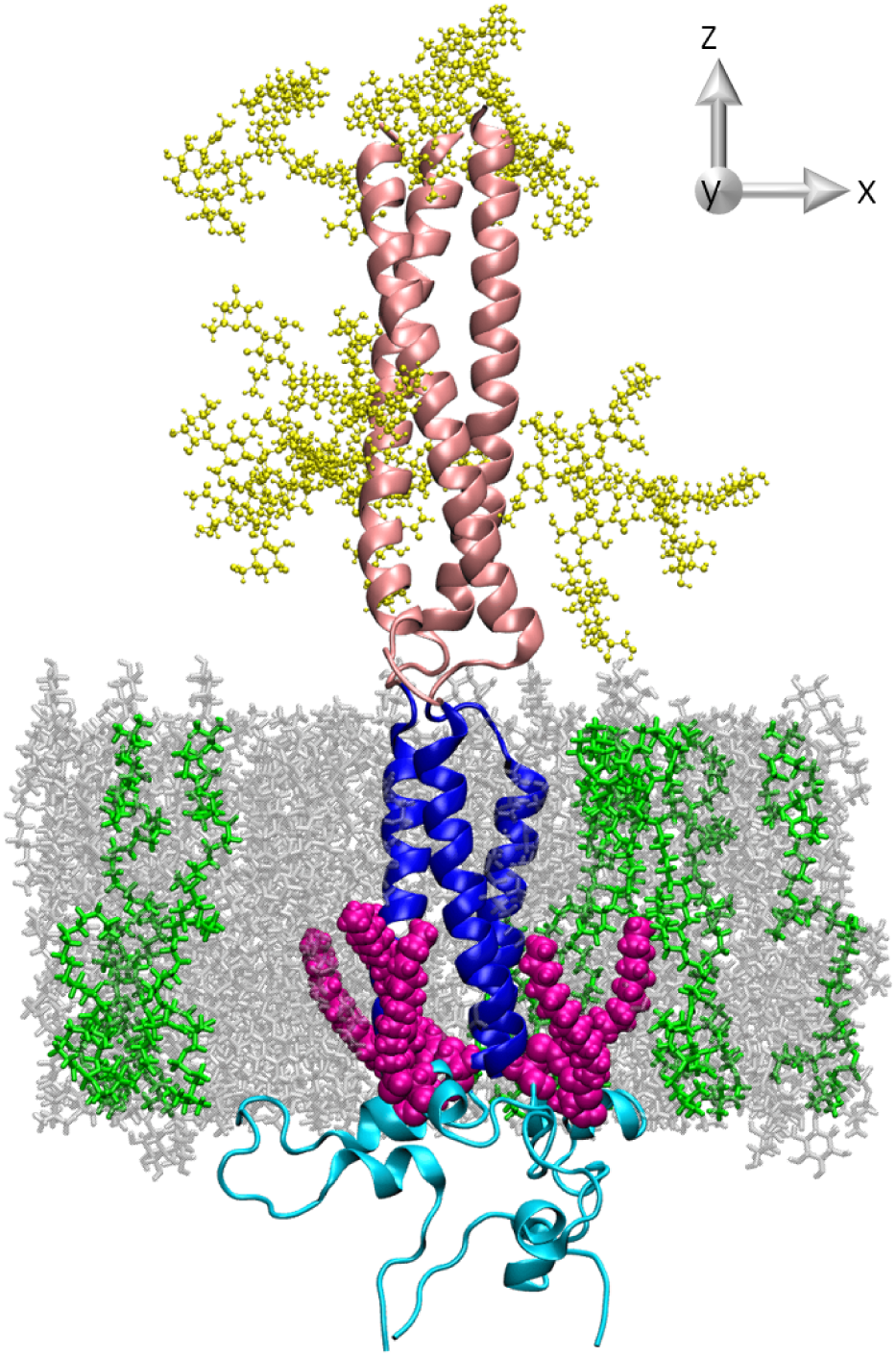
The simulation setup with 25% POPC oxidation. The spike protein segment (residues 1168 − 1273) is shown schematically with the HR2 domain in pink, the TM helix in blue, and the CT domain in cyan. Palmitic acids are displayed in magenta using the van der Waals representation, glycans in yellow using the CPK style, and PoxnoPC lipids in green using a licorice representation. All molecular graphics were generated using VMD [35].

From this model, we extracted residues 1168 − 1273, corresponding to the C-terminal HR2 segment (shown in pink), TM region (blue), and CT domain (cyan). The resulting trimeric fragment was then oriented along the membrane normal (*z*-axis) using the Orientations of Proteins in Membranes (OPM) database [33] prior to membrane embedding. The initial membrane systems were constructed using the Membrane Builder module of the CHARMM-GUI input generator [34]. The lipid composition of the non-oxidized reference membrane followed the ERGIC-like model used by Casalino et al. [23], consisting of approximately POPC (47%), POPE (20%), cholesterol (CHL) (15%), POPI (11%), and POPS (7%). To investigate the effects of lipid peroxidation, we generated five membrane systems containing 0%, 25%, 50%, 75%, and 100% oxidized POPC by replacing a corresponding fraction of POPC molecules with their aldehyde-containing oxidation product PoxnoPC using CHARMM-GUI’s oxidized lipid library. Each viral membrane consisted of 198 phospholipids, which were evenly distributed between the leaflets. The membrane was aligned with the *xy*-plane, with its normal oriented along the *z*-axis. The exact POPC and PoxnoPC compositions for each oxidation level are listed in Table 1.

**Table 1:** POPC and PoxnoPC composition at each oxidation level. The last column shows the percentage of oxidized lipids relative to all PO-type glycerophospholipids (POPC + POPE + POPI + POPS = 85%).

| Oxidation level (%) | POPC (%) | PoxnoPC (%) | Oxidized PO lipids (%) |
| --- | --- | --- | --- |
| 0 | 47.0 | 0.0 | 0.0 |
| 25 | 35.2 | 11.8 | 13.9 |
| 50 | 23.5 | 23.5 | 27.6 |
| 75 | 11.8 | 35.2 | 41.4 |
| 100 | 0.0 | 47.0 | 55.3 |

**Table 2:** Area per lipid (APL, Å^2^) of individual lipid species as a function of oxidation level. Values were calculated over the final 50 ns of each trajectory, averaged over all analyzed frames for each replica. Final values represent mean ± standard deviation from three independent replicas.

|  | POPC | PoxnoPC | POPE | CHL | POPI | POPS |
| --- | --- | --- | --- | --- | --- | --- |
| 0% | $55.7 \pm 0.2$ | | $50.5 \pm 0.8$ | $38.2 \pm 0.4$ | $55.2 \pm 0.3$ | $53 \pm 1$ |
| 25% | $57.7 \pm 0.6$ | $59 \pm 1$ | $52.2 \pm 0.3$ | $38.2 \pm 0.6$ | $58 \pm 1$ | $55 \pm 1$ |
| 50% | $58.2 \pm 0.9$ | $61.4 \pm 0.2$ | $51.3 \pm 0.6$ | $41.5 \pm 0.8$ | $57.7 \pm 0.5$ | $54.4 \pm 0.5$ |
| 75% | $56.2 \pm 0.5$ | $61.6 \pm 0.6$ | $52 \pm 1$ | $40 \pm 1$ | $57.2 \pm 0.5$ | $54.8 \pm 0.7$ |
| 100% | | $61.8 \pm 0.2$ | $54.1 \pm 0.8$ | $40.1 \pm 0.4$ | $57 \pm 1$ | $58 \pm 1$ |

Next, for all systems, a palmitoyl thioester was attached to cysteine residues 1236 and 1241 in each chain of the S protein [23]. The protein was glycosylated at residues 1173 and 1194 using the same glycan compositions as reported by Casalino et al. [23]. The system was placed in a rectangular simulation box, with dimensions ranging from 8.3 × 8.3 × 18.4 nm^3^ to 8.3 × 8.3 × 19.2 nm^3^ in the *x*, *y* and *z* directions. Additionally, approximately 30 000 water molecules were inserted into the box. The TIP3P water model was used for all systems. Chloride and sodium ions were added to neutralize all simulation systems. The final systems contained between 118 000 and 122 000 atoms in total. After equilibration, the S protein segment corresponding to HR2 (pink region in Fig. 1), including its attached glycans, was truncated near the pulling site to ensure that the cytoplasmic tail remained inside the simulation box during the pulling simulations used for umbrella sampling (US). Only residues 1210 − 1273 were included and each N-terminus was capped by adding two additional hydrogen atoms. Moreover, to provide space in the *z*-direction for pulling simulations, the system was first moved 3 nm to the negative *z*-direction and equilibrated for 0.1 ns to allow all the water molecules to fill the box. The box was then extended 22 nm in the positive *z*-direction, and approximately 45,000 water molecules were inserted. Finally, these modified systems were neutralized with chloride ions and contained between 250,000 and 254,000 atoms.

### 2.2. Computational details

All MD simulations were performed using GROMACS 2023.3 software [36] with the CHARMM36 all-atom force field [37, 38]. Periodic boundary conditions were applied in all directions. For both van der Waals and Coulombic interactions, a cut-off distance of 1.2 nm was applied, and for longrange interactions, the fast smooth Particle-Mesh Ewald (SPME) electrostatics [39] was used. The LINCS algorithm [40] was used for the bond constraints. Energy minimization was performed using the steepest descent algorithm with the Verlet cutoff scheme and a force tolerance value of 1000 kJ mol^−1^ nm^−1^.

After minimization of each system, three replicas were generated with different random initial velocities. The first step of equilibration in the *NV T* ensemble was performed for 2 ns under temperature control using the Berendsen thermostat [41], with a reference temperature of 310 K. The time constant for temperature coupling was 1.0 ps. After that, all systems were relaxed in the *NPT* ensemble for 2 ns using the Berendsen thermostat and barostat [41], corresponding to a semiisotropic setup. In these equilibration steps, a reference temperature of 310 K, a temperature time constant of 1.0 ps, a reference pressure of 1.0 bar, compressibility of 4.5×10^−5^ bar^−1^ and a pressure time constant of 5.0 ps were applied. The dynamics production simulations were performed in the *NPT* ensemble for 150 ns using the Nosé-Hoover thermostat [42–44] and the Parrinello-Rahman barostat [45] with the same parameters as for the Berendsen thermostat and barostat. Protein equilibration was assessed using the C*_α_* root-mean-square deviation (RMSD) of the TM–CT region, and membrane-property analyses were conducted only after the RMSD reached a stable plateau. In all systems, convergence occurred well before the final 50 ns of the production trajectories used in structural analysis. RMSD profiles for all oxidation levels and replicas are provided as Supporting Information.

The same equilibration protocol was used for the modified systems, which included the cut protein in an extended box. Due to the increased number of atoms, each equilibration step lasted 2 ns. A 40 ns production run was then performed for these systems using the same parameters as for the original systems. The final structures from these production trajectories were used as starting configurations for the constant-velocity steered MD (CVSMD) simulations. In these, the S protein tail was pulled from the viral membrane along the positive *z*-axis over a period of 6000 ps, where snapshots were saved every 10 ps. The pulling force was applied to the center of mass (COM) of the protein, and the membrane was kept as a reference group. A spring constant of 1500 kJ mol^−1^ nm^−2^ and a pulling rate of 0.005 nm/ps were applied in all CVSMD simulations. A position restraint of 1000 kJ mol^−1^ nm^−2^ was applied in the *z*-direction for all phosphorus atoms in the membrane to prevent any parts of the membrane from being pulled along with the protein. We note that position restraints on membrane phosphorus atoms during the CVSMD simulations, while necessary to prevent lipid co-translation, suppress local membrane deformations. However, these restraints were removed during the umbrella sampling calculations used to construct the PMF, allowing the membrane to relax within each window. As a result, any influence on the free-energy profiles is expected to arise primarily from the initial configurations rather than from direct bias in the sampling. While this may affect the absolute magnitude of the PMF to some extent, the consistent application of the same protocol across all oxidation levels ensures that the relative freeenergy differences, which constitute our central conclusion, remain robust.

The stability of PMF profiles with increasing sampling time further supports adequate relaxation of the dominant membrane modes within the production windows. The frames of the CVSMD trajectories were then extracted to serve as starting configurations for the US simulations. Trajectories were analyzed up to a COM distance of 15 nm. The windows were spaced at 0.1 nm intervals along the reaction coordinate, which was defined as the COM distance between the S protein tail and the viral membrane. Accordingly, frames where this COM distance differed by approximately 0.1 nm were selected. This spacing yielded 128 − 142 windows per replica, resulting in approximately 400 windows per oxidation level.

The position restraints applied for membrane phosphorus atoms during SMD simulations were removed for US simulations. For each window, US simulation, consisting of 5 ns of equilibration and 30 ns of sampling, was performed in the *NPT* ensemble by applying the Nosé-Hoover thermostat with a reference temperature of 310 K and a temperature time constant of 1.0 ps. A harmonic bias potential was applied between the centers of mass of protein and membrane along the *z*-axis, with a force constant of 1000 kJ mol^−1^ nm^−2^. This potential restrained the COM distance between protein and membrane to remain close to a window-specific reference coordinate.

Finally, potential of mean force (PMF) profiles were obtained from the US simulations using the Weighted Histogram Analysis Method (WHAM) [46–48]. The final PMF for each system was obtained by averaging over 3 individual PMFs (using 3 trajectories, see above). Thus, to obtain the final PMFs, approximately 3 × 130 ≈ 390 US windows were used for each system. For error estimation, a bootstrap method with 100 bootstraps was used. Convergence of the PMF was assessed by comparing cumulative free-energy profiles reconstructed from increasing portions of the sampling trajectory (e.g., the first 10, 20, and 30 ns of production data for each window). In addition, histogram overlap between adjacent windows was inspected to verify adequate sampling continuity along the reaction coordinate. The final PMF profiles were considered converged when the overall profile shape and barrier height remained stable with increasing sampling time and sufficient overlap was observed between neighboring windows. Windows showing insufficient overlap were extended until stable PMF reconstruction was obtained.

### 2.3. Membrane property analysis

To quantify the structural effects of POPC oxidation on the viral membrane, we evaluated lipid tail order parameters, number density distributions, and lateral lipid clustering at all oxidation levels. These quantities were calculated by averaging over three independent replicas for each oxidation level, using the final 50 ns. The definitions of the lipid structural regions used in these analyses (headgroup, glycerol–ester linkage, and acyl chain segments) are illustrated in Figure 2.

**Figure 2:**
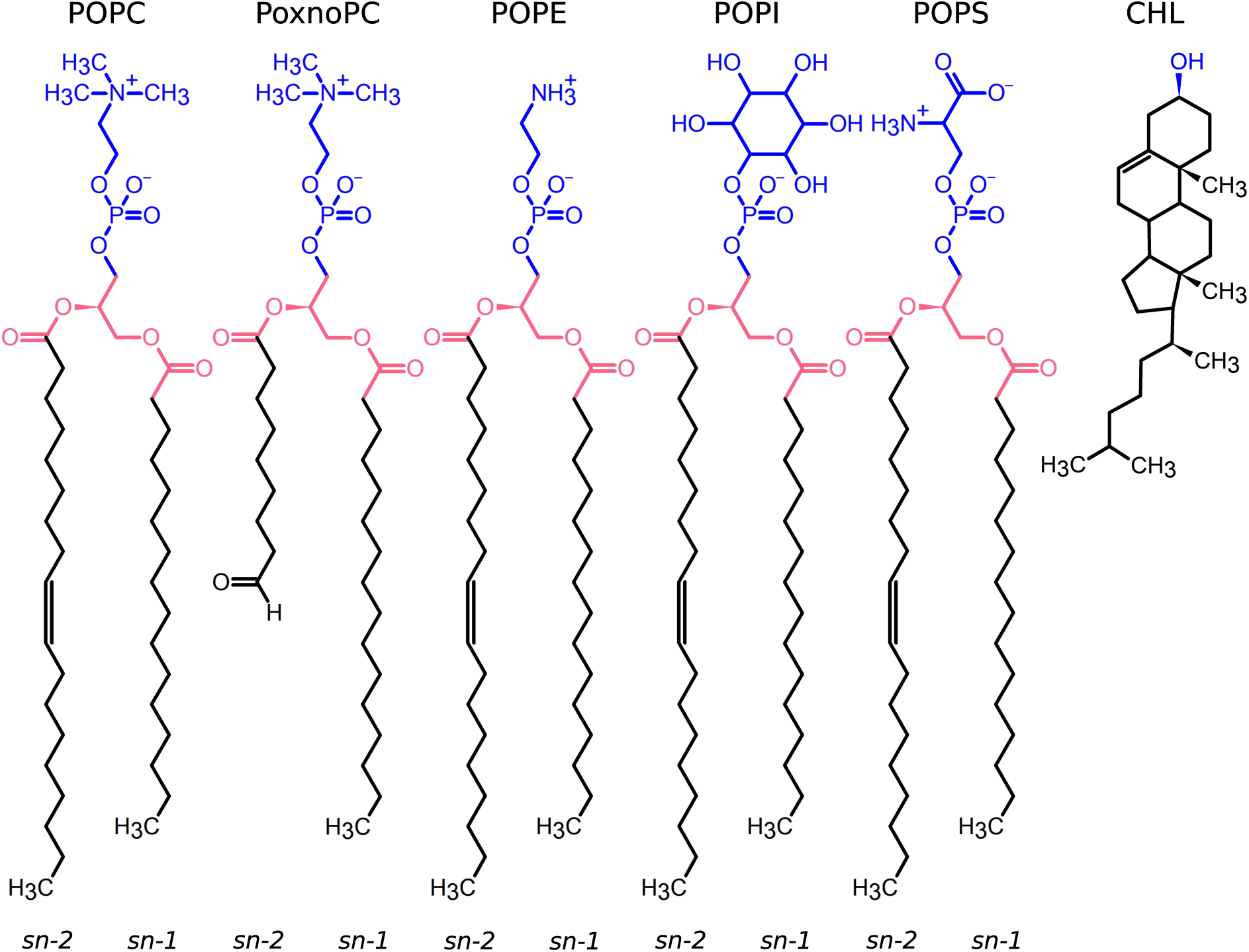
Chemical structures of membrane phospholipids and cholesterol (CHL) used in this study. The headgroup, the glycerol–ester linkage, and acyl chain regions of phospholipids are highlighted in blue, pink and black colors, respectively. The head group of CHL is shown in blue. The positions of the *sn*-1 and *sn*-2 acyl chains are indicated, showing that PoxnoPC results from oxidation of the *sn*-2 chain. While POPC, PoxnoPC and POPE are zwitterionic under physiological conditions, POPI and POPS are anionic with a net charge of -1.

### Lipid tail order parameters

The conformational order of lipid acyl chains was quantified by the deuterium order parameter *S*_CD_, defined as [49]

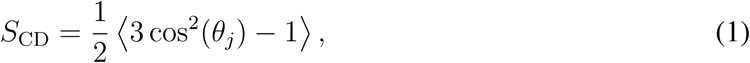

where *θ_j_* is the angle between the membrane normal and a specific C-H bond vector, that is, the vector between adjacent carbon atoms (C*_n_*_−1_– C*_n_*), corresponding to the experimental C–D bond direction, and the membrane normal. Values of *S*_CD_ = 1 correspond to lipid chains fully aligned with the membrane normal, whereas *S*_CD_ = −1*/*2 corresponds to chains lying perpendicular to it [7]. A reduction in the magnitude of *S*_CD_ reflects increased chain disorder and membrane fluidity [6].

Order parameters were computed separately for the *sn*-1 and *sn*-2 acyl chains using to distinguish oxidation-dependent effects on the two chains. The order parameters were computed for all lipid species (POPC, PoxnoPC, POPE, POPI and POPS), and the reported values are ensemble averages profiles over three replicas and both leaflets. Because oxidation of POPC disrupts a portion of the *sn*-2 acyl chain (i.e., atoms beyond C7 are removed in PoxnoPC), direct comparison of the full *sn*-2 chain between oxidized and non-oxidized lipids is not meaningful. Therefore, for *sn*-2 analysis we quantified order parameters only for the C1 – C7 segment, which is structurally conserved across all glycerophospholipids. In contrast, the *sn*-1 chains remain intact in all systems and their order parameters were calculated throughout the chain length. This approach enables consistent quantification of oxidation-induced disorder within the common acyl-chain region without bias from chain truncation in PoxnoPC. The Lipid-specific order parameter profiles for each phospholipid type are provided separately in the SI.

### Membrane density profiles

For each lipid species, atoms were grouped into three structural segments: (1) the headgroup (choline and phosphate atoms); (2) the glycerol–ester linkage, defined by the glycerol backbone, carbonyl and ester linkage atoms connecting the headgroup and the acyl chains; and (3) the tail region, defined as all remaining aliphatic carbons of the *sn*-1 and *sn*-2 acyl chains (Fig.2).

The spatial distribution of these atomic groups along the membrane normal was characterized by computing number density profiles. Density profiles were averaged over all production trajectories and compared across oxidation levels to asses changes in bilayer thickness, interfacial sharpness, and hydrophobic-core organization. Number density profiles along the membrane normal (*z*-axis) were computed for all systems. For an atomic group with ⟨*N* (*z*)⟩ atoms within a slab of thickness Δ*z* at position *z*, the number density *ρ*(*z*) is given as

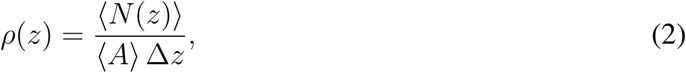

where ⟨*A*⟩ = ⟨*L_x_L_y_*⟩ is the instantaneous cross-sectional area of the simulation box, averaged over the analyzed frames. Defining the bilayer center as *z* = 0, density profiles were symmetrized with respect to the membrane midplane, yielding identical leaflet contributions. For quantitative comparison across oxidation levels, each profile was normalized by its total integrated density:

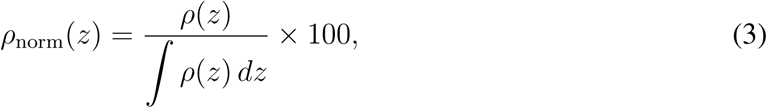

yielding the relative number density (% per nm). Normalized density profiles were averaged over all production frames from three independent replicas for each oxidation level.

### Area per lipid and bilayer thickness

Membrane packing was characterized by calculating the area per lipid (APL) using the `FATSLiM` toolkit [50]. In this approach, APL is estimated for each lipid from its accessible area obtained by a local Voronoi tessellation of headgroup positions projected onto the membrane plane. For phospholipids, the phosphorus atom (P) was used as the representative headgroup coordinate, whereas for CHL the oxygen atom (O) was used. Because the simulated systems contain an embedded protein, protein atoms were included in the analysis to account for the protein-excluded membrane area and to avoid artificial enlargement of lipid areas near the protein surface.

For a given lipid species *L*, the time-averaged APL for each replica, calculated over the final 50 ns of the trajectory, was computed as:

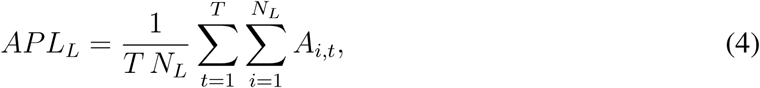

where *A_i,t_*is the instantaneous area associated with lipid *i* at frame *t*, *N_L_*is the number of lipids of type *L*, and *T* is the total number of analyzed frames.

To visualize the spatial distribution of membrane packing, two-dimensional APL maps were generated from the `FATSLiM` output for each oxidation level. For each frame, per-lipid APL values and headgroup coordinates were extracted and assigned to the upper or lower leaflet. The membrane plane was divided into a regular 50 × 50 grid based on the box dimensions, and the APL values of lipids falling within each bin were averaged over all lipids in that bin and over all analyzed frames. Separate maps were constructed for the upper and lower leaflets. To facilitate visual comparison between oxidation levels, a common color scale was used for all maps, centered on the global mean APL of the non-oxidized bilayer and displayed symmetrically within a fixed range of ±50% around this reference value.

In addition, membrane thickness was calculated using `FATSLiM`. In this method, local interleaflet distances are estimated from neighborhood-averaged positions of lipids in the two opposing leaflets, and the membrane thickness for each frame is reported as the membrane-averaged value over all analyzed lipids. The time-averaged membrane thickness for each replica, calculated over the final 50 ns of the trajectory, was computed as

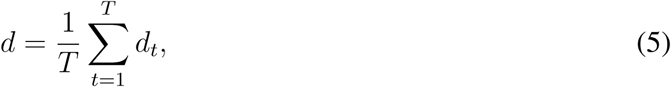

where *d_t_*is the membrane thickness reported by `FATSLiM` at frame *t* and *T* is the number of analyzed frames. For both APL and thickness, final values are reported as the replica-averaged mean and standard deviation.

### Lipid clustering

Lateral lipid clustering was analyzed to quantify oxidation-dependent changes in membrane organization. A two-step procedure was used. First, radial distribution functions (RDFs) were computed between P atoms for phospholipids (POPC, POPE, PoxnoPC), O atoms for CHL, and P–O for the mixed pairs. For each pair type, the cutoff distance defining cluster membership was chosen to be the first minimum of the corresponding RDF. Interactions with CHL (POPC–CHL, PoxnoPC–CHL, and POPC–PoxnoPC–CHL mixtures) required a larger cutoff of 0.90 nm due to the bulky sterol structure of CHL. For POPE–POPE interactions, a cutoff of 0.90 nm was obtained. RDFs for POPS and POPI exhibited substantially larger first minima (2.0 nm and 1.4 nm, respectively) due to extended electrostatic interactions arising from their negatively charged headgroups; their low abundance, however, precluded separate clustering analysis.. Therefore, clustering metrics reported here focus on the PC lipids and their oxidation-dependent interactions with CHL. In the clustering process, each lipid was assigned to a cluster if its reference atom (P for phospholipids, O for CHL) lay within the RDF-derived cutoff of at least one other lipid belonging to the same cluster. The resulting histograms provide the average number of clusters per frame, *N*_clust_(*s*), for each cluster size *s*. For a lipid subset containing *N*_tot_ molecules (e.g., POPC+PoxnoPC or POPC+PoxnoPC+CHL), the fraction of lipids participating in clusters of size *s* was computed as

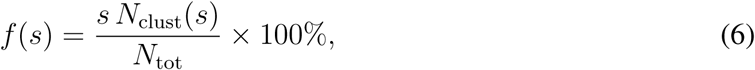

where the numerator gives the average number of lipids belonging to clusters of size *s*. This calculation was applied to each replica separately; final values are reported as the replica-averaged mean and standard deviation. To reflect distinct structural regimes, cluster sizes were grouped into three classes: proto-clusters (3 − 4 lipids), small clusters (5 − 10 lipids), and large clusters (*s >* 10). Reported percentages therefore represent the proportion of phospholipids involved in clusters of each class, averaged over time and over three independent simulations.

## 3. Results and Discussion

### 3.1. Free-energy of spike–membrane detachment

To quantify how lipid peroxidation alters the stability of the SARS-CoV-2 spike within the viral membrane, PMF profiles were obtained by extracting the TM helix and CT domain along the membrane normal using US simulation (Fig. 3). The pulling direction was chosen along the membrane normal toward the aqueous phase, so as to represent the biologically relevant pathway for protein detachment from the bilayer.

**Figure 3:**
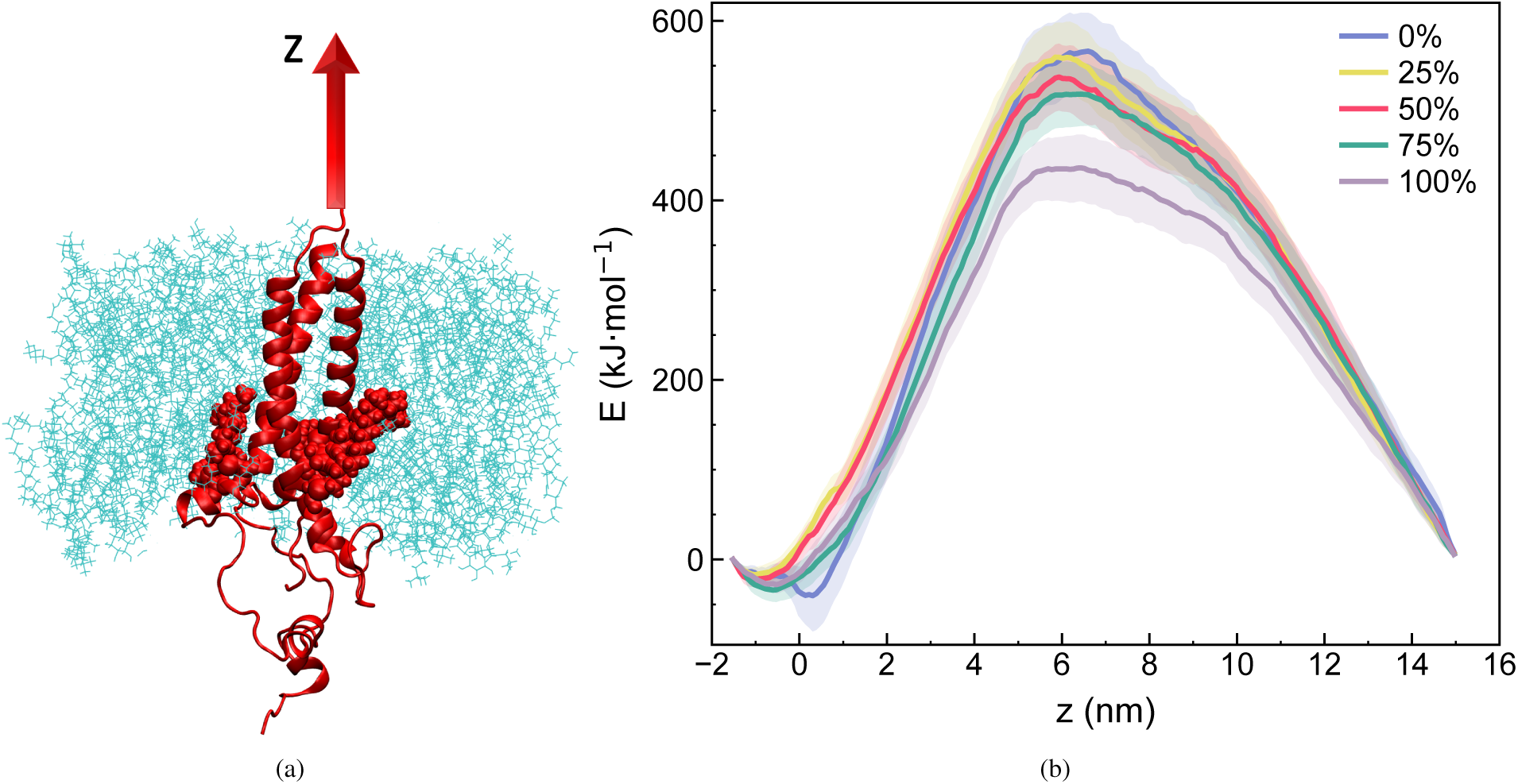
PMF profiles for extraction of the spike TM helix together with its CT domain from the viral membrane at different POPC oxidation levels. The free energy is plotted as a function of the center-of-mass (COM) separation between the TM+CT region and the membrane along the membrane normal (*z* axis). (a) Schematic representation of the system and reaction coordinate used in the PMF calculations. The spike TM+CT region is shown in red (new cartoon representation), with palmitoylated cysteine residues highlighted in red van der Waals representation, while the membrane is shown in cyan line representation. The red arrow indicates extraction of the TM+CT region from the membrane along the *z*-direction. (b) PMF curves including bootstrap standard deviations (error bars), computed from three independent replicas at each oxidation level, demonstrating a statistically robust, oxidation-dependent weakening of spike–membrane coupling along the reaction coordinate defined in panel (a). Curves correspond to 0% (blue), 25% (yellow), 50% (red), 75% (green), and 100% (lilac) PoxnoPC substitution.

In the native membrane (0% PoxnoPC), the anchoring free-energy barrier was 606±39 kJ mol^−1^, reflecting strong hydrophobic engagement of the TM helix with the bilayer. Increasing the level of POPC oxidation resulted in a decreasing trend in the detachment barriers, although the differences at intermediate oxidation levels were not statistically significant given the sampling uncertainties:

576 ± 39 kJ mol^−1^ at 25% PoxnoPC, 560 ± 38 kJ mol^−1^ at 50%, and 552 ± 38 kJ mol^−1^ at 75%. These modest differences suggest that low-to-moderate oxidation softens the membrane and weakens spike anchoring, but does not fully compromise membrane attachment.

In the fully oxidized POPC case (100% PoxnoPC), the barrier was significantly reduced to 464 ± 38 kJ mol^−1^ – a reduction of ≈ 140 kJ mol^−1^ (≈ 23%) relative to the native system.

Using propagated uncertainty of the difference gives significance of 2.6 standard deviations showing the difference in the barrier heights is statistically robust. Importantly, this condition corresponds to oxidation of only ≈ 55% of all PO-type phospholipids in the bilayer, because POPC constitutes 47% of the total lipid composition (Table 1). Thus, destabilization of spike anchoring does not require complete membrane oxidation, as oxidation of roughly half of the glycerophospholipids is sufficient to produce a measurable reduction in TM–bilayer coupling strength.

We note, however, that even the reduced barrier of 464 kJ mol^−1^ corresponds to approximately 180*k*_B_*T* at 310 K, which remains exceedingly large compared to thermal energy. While position restraints on membrane phosphorus atoms were applied during the CVSMD stage to prevent lipid co-translation, these restraints were removed during the umbrella sampling simulations used to construct the PMF, allowing the membrane to relax within each sampling window. As a result, the free-energy profiles are not directly biased by the restraints; however, the restrained CVSMD pathway may influence the initial configurations and thus the absolute magnitude of the barrier to some extent, such that the reported values may represent an upper bound of the true free-energy barrier. Importantly, because the same protocol was applied across all oxidation levels, the relative reduction in the barrier remains robust.

Spontaneous detachment of the spike TM region by thermal fluctuations alone is therefore not expected on any biologically relevant timescale, even in fully oxidized membranes. The significance of the 23% reduction should instead be interpreted in the context of cooperative processes such as mechanical forces during membrane fusion, immune-mediated disruption, or simultaneous oxidative damage to the protein that may act synergistically with membrane weakening to compromise viral function.

Together, these PMF results demonstrate a statistically significant effect of extensive peroxidation on spike–membrane coupling: while intermediate oxidation levels (25% 75% POPC replacement) do not produce statistically resolvable changes in anchoring strength, full POPC oxidation (approx. 55% of all PO-type lipids) reduces the extraction barrier by about 23%, consistent with oxidative damage contributing to antiviral mechanisms against SARS-CoV-2. Whether this reduction is sufficient to compromise spike function *in vivo* likely depends on the coupling between membrane weakening and other disruptive processes.

### 3.2. Oxidation reduces acyl-chain order in all membrane phospholipids

To assess how POPC oxidation alters bilayer packing, deuterium order parameters (*S*_CD_) were calculated for the acyl chains of each major phospholipid species (POPC, PoxnoPC, POPE, POPI, and POPS) in the viral membrane (Figure 4). This species-resolved analysis allows for a direct assessment of how oxidation perturbs both local and collective acyl chain order. Chain-resolved order parameters for each individual lipid type are provided in the SI.

**Figure 4:**
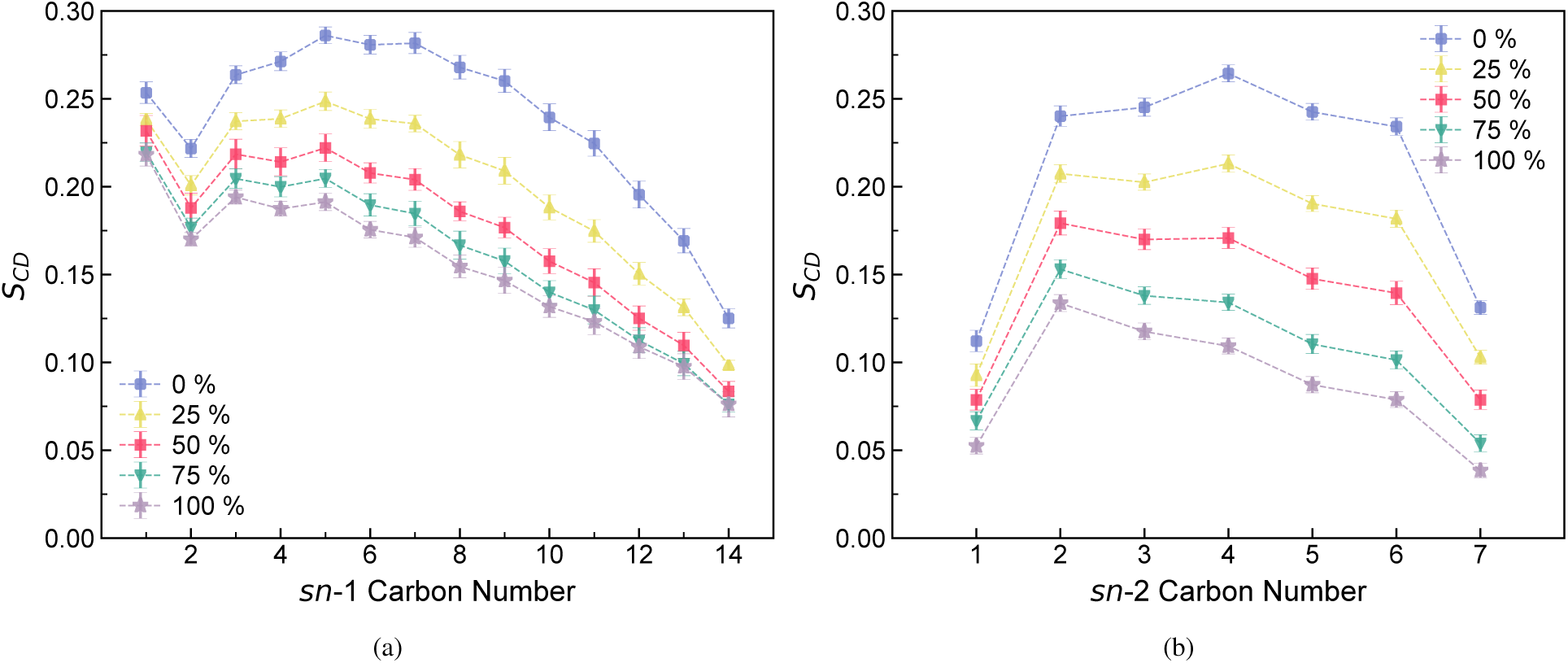
Deuterium order parameters (*S*_CD_) averaged over all glycerophospholipids for the *sn*-1 (a) and *sn*-2 (b) chains at different oxidation levels (0% blue, 25% yellow, 50% red, 75% green, 100% lilac). Oxidation progressively increases acyl-chain disorder, with a stronger effect on *sn*-2 due to the structural defect introduced by PoxnoPC. *sn*-2 order parameters reflect only the conserved C1 – C7 segment, because oxidation shortens the PoxnoPC *sn*-2 chain.

In the non-oxidized membrane, both *sn*-1 and *sn*-2 chains display high *S*_CD_ values in the bilayer core, reflecting tight packing. As the level of PoxnoPC increases, *S*_CD_ decreases systematically for both chains, demonstrating that oxidation weakens hydrophobic ordering across the bilayer. This disordering is not limited to the oxidized lipid species: even unmodified lipids (POPE, POPI, POPS) exhibit reduced order in oxidized systems, indicating that structural defects introduced by PoxnoPC propagate into their local environment. We note that the direction and magnitude of oxidation-induced ordering changes depend sensitively on membrane composition and oxidation regime. In homogeneous POPC-based membranes, Bagheri et al. [24] reported that introducing PoxnoPC at low oxidized fractions (up to ≈30%) produces only modest changes and, in some cases, a slight increase in the order of the remaining POPC chains. In contrast, the present ERGIC-like viral envelope is compositionally heterogeneous, contains CHL, and is examined over a substantially broader oxidation range (up to complete POPC substitution, corresponding to ≈ 55% oxidation of all PO-type glycerophospholipids). In this complex environment, oxidized-lipid packing defects and altered lateral organization dominate the collective response, yielding a net reduction in acylchain order across lipid species. A key feature of this global disordering is that the reduction is consistently more pronounced in *sn*-2 chains than in *sn*-1, especially at high oxidation levels. This asymmetry originates from the truncated and kinked *sn*-2 chain of PoxnoPC, which disrupts local packing and increases free volume within the membrane interior. At the highest oxidation level (100% PoxnoPC, corresponding to ≈ 55% total PO-lipid oxidation), the average *S*_CD_ decreases by approximately 30 − 35% compared to the native system, indicating a transition toward a more disordered and fluid bilayer. Thus, POPC peroxidation not only alters the conformational order of the oxidized lipids themselves, but also reduces acyl-chain ordering across the entire membrane, which is consistent with the weakened spike anchoring observed in the PMF results. Establishing a causal link would require decomposition of the free-energy contributions (e.g., via separate PMFs with and without palmitoylation) or lateral pressure profile analysis. These tasks are beyond the scope of the current study.

#### 3.2.1. Area per lipid and bilayer thickness

The APL values for each lipid species as a function of oxidation level are summarized in Table 2.

The analysis reveals a composition-dependent response to lipid oxidation. As the fraction of oxidized lipid increased from 0% to 100%, the APL of PoxnoPC remained consistently larger than that of POPC at comparable oxidation levels. At 25% oxidation, the APL values of POPC and PoxnoPC were 57.7 ±0.6 and 59 ±1 Å^2^, respectively, and this difference became more pronounced at 50% oxidation, where POPC and PoxnoPC exhibited APL values of 58.2±0.9 and 61.4±0.2 Å^2^. At 75% oxidation, PoxnoPC remained substantially expanded (61.6 ± 0.6 Å^2^), whereas POPC decreased slightly to 56.2±0.5 Å^2^. In the fully oxidized membrane, PoxnoPC reached 61.8±0.2 Å^2^. Overall, these results indicate that oxidation increases the lateral area occupied by the oxidized lipid and promotes membrane expansion, although the response of POPC is non-monotonic and suggests a more complex reorganization of the mixed bilayer environment.

The other lipid species showed smaller but still notable composition-dependent changes. POPE remained comparatively compact throughout the oxidation series, varying from 50.5±0.8 Å^2^ at 0% oxidation to 54.1 ± 0.8 Å^2^ at 100% oxidation. POPI and POPS also exhibited moderate increases, with POPI changing from 55.2 ± 0.3 to 57 ± 1 Å^2^ and POPS from 53 ± 1 to 58 ± 1 Å^2^. CHL showed the smallest absolute area among all components, ranging from 38.2 ± 0.4 to 41.5 ± 0.8 Å^2^, consistent with its well-known condensing role in mixed membranes.

The observed APL values are in reasonable agreement with previous simulation studies, while also reflecting differences in membrane composition, oxidation state, force field, and temperature. For pure or non-oxidized POPC bilayers, literature values generally fall in the range of 60 − 65 Å^2^ [5, 24, 51–53], although the exact value depends on the simulation conditions. In the heterogeneous-membrane study of Kumar and Sastry [51], APL values of 64.81 Å^2^ and 57.02 Å^2^ were reported for pure POPC and POPE bilayers, respectively, at 310.15 K using the CHARMM36 force field. When these lipids were mixed with other phospholipid components, the APL values changed depending on membrane composition.

In the study of Kumar and Sastry [51], mixed membranes with increasing compositional complexity showed a reduction in the average membrane APL, particularly in systems containing CHL and sphingomyelin (PSM), where the average APL decreased to approximately 45−49 Å^2^, indicating strong membrane condensation. Their component-resolved values for the mixed POPC/POPE membrane were 60.21 and 59.30 Å^2^ for POPC and POPE, respectively, whereas in more complex bilayers containing POPC, POPE, POPI, POPS, PSM, and CHL these values decreased to 56.60 and 55.69 Å^2^. The authors further showed that changes in the relative lipid composition also affect both the component-resolved APL and the overall membrane APL. Thus, their results demonstrate that increasing membrane heterogeneity, and especially the presence of condensing lipids such as CHL and PSM, reduces the average membrane area per lipid. The APL of CHL obtained here (38 − 42 Å^2^) is larger than the component values reported in the Kumar and Sastry models (≈ 27 − 29 Å^2^), which likely reflects methodological differences in component-area definitions, as well as differences in membrane composition and local environment. Importantly, despite these quantitative differences, both datasets support the same qualitative conclusion: cholesterol contributes to tighter membrane packing, whereas lipids associated with more disordered or expanded local environments occupy larger lateral area. The oxidation trend observed here is also consistent with previous reports on oxidized phospholipid bilayers. In the literature values collected above, PoxnoPC-containing membranes generally show increased APL relative to non-oxidized POPC-rich systems, and the oxidized lipid itself often occupies a larger lateral area than POPC. For example, previous GROMOS-based simulations [5, 24] reported PoxnoPC APL values near 59.4 − 65.0 Å^2^ depending on oxidation level, while POPC in mixed POPC/PoxnoPC systems remained lower, around 59.0 − 61.0 Å^2^. The present results fall well within this range, with PoxnoPC increasing from 59 ± 1 to 61.8 ± 0.2 Å^2^ across the oxidation series, supporting the interpretation that oxidation of the *sn*-2 chain promotes lateral expansion of the oxidized lipid. At the same time, the non-monotonic behavior of POPC suggests that the membrane response is not a simple uniform expansion, but rather a redistribution of local packing constraints among oxidized and non-oxidized lipids, cholesterol, and the protein surface.

Overall, the APL results indicate that membrane oxidation primarily affects local packing by increasing the lateral area of the oxidized lipid species, while the remaining lipid components respond more moderately. In comparison with previous homogeneous and heterogeneous bilayer studies, the present values are consistent with the expected balance between two competing effects: oxidation-driven membrane expansion and cholesterol-/protein-associated condensation. This interplay likely underlies the non-linear changes observed for POPC and the relatively restrained response of POPE, POPI, and POPS.

The upper leaflet-resolved two-dimensional APL maps show that membrane packing is spatially heterogeneous at all oxidation levels (Figure 5), as expected for a bilayer with a chemically heterogeneous lipid composition. Oxidation therefore does not create heterogeneity per se, but rather modifies the pre-existing lateral packing landscape. In particular, the upper-leaflet maps suggest that increasing oxidation enhances the extent of high-APL regions at the protein boundary, visible as red patches surrounding the protein footprint. This trend suggests a local expansion of lipid packing in the immediate vicinity of the protein upon oxidation. At the same time, low-APL regions remain distributed in the surrounding membrane, indicating that the oxidation-dependent response is spatially non-uniform and strongly influenced by the local membrane environment. A similar qualitative trend was also observed in the lower leaflet.

**Figure 5:**
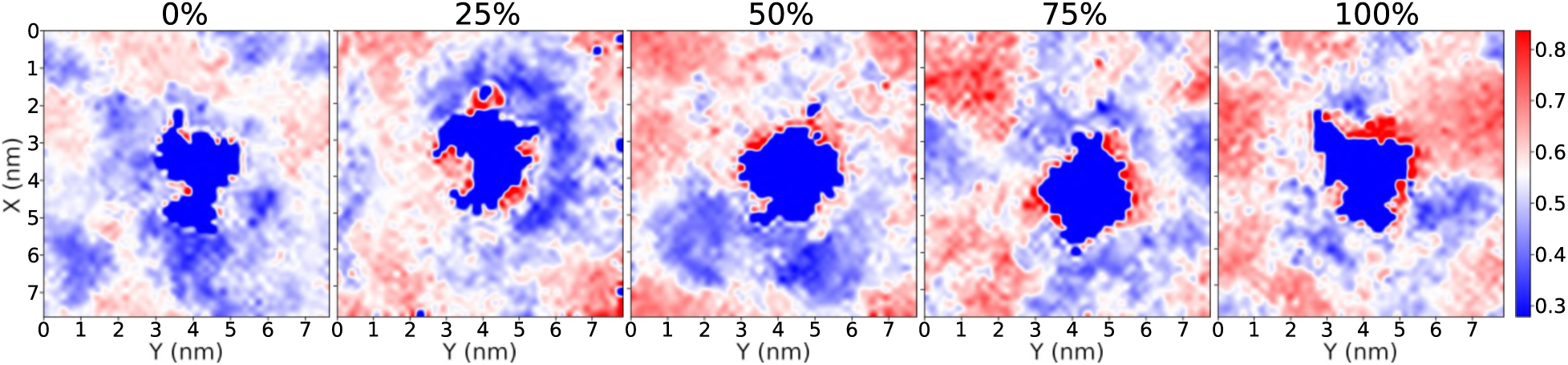
Leaflet-resolved two-dimensional maps of the local mean APL at different oxidation levels. The maps shown here correspond to the upper leaflet; a similar qualitative trend was also observed for the lower leaflet. Maps were generated from the `FATSLiM` per-lipid output by assigning lipid coordinates in the membrane plane to a 50 × 50 grid and averaging the APL values within each bin over all analyzed frames. A common color scale, centered on the global mean APL of the non-oxidized bilayer, was used to facilitate comparison between oxidation levels. The central dark-blue region corresponds to the protein footprint, where no lipid APL values are present.

The membrane thickness as a function of oxidation level is shown in Figure 6. A clear monotonic decrease in thickness is observed with increasing oxidation, dropping from approximately 4.2 nm in the non-oxidized system to ≈ 3.3 nm in the fully oxidized bilayer. This trend indicates that lipid oxidation progressively thins the membrane.

**Figure 6:**
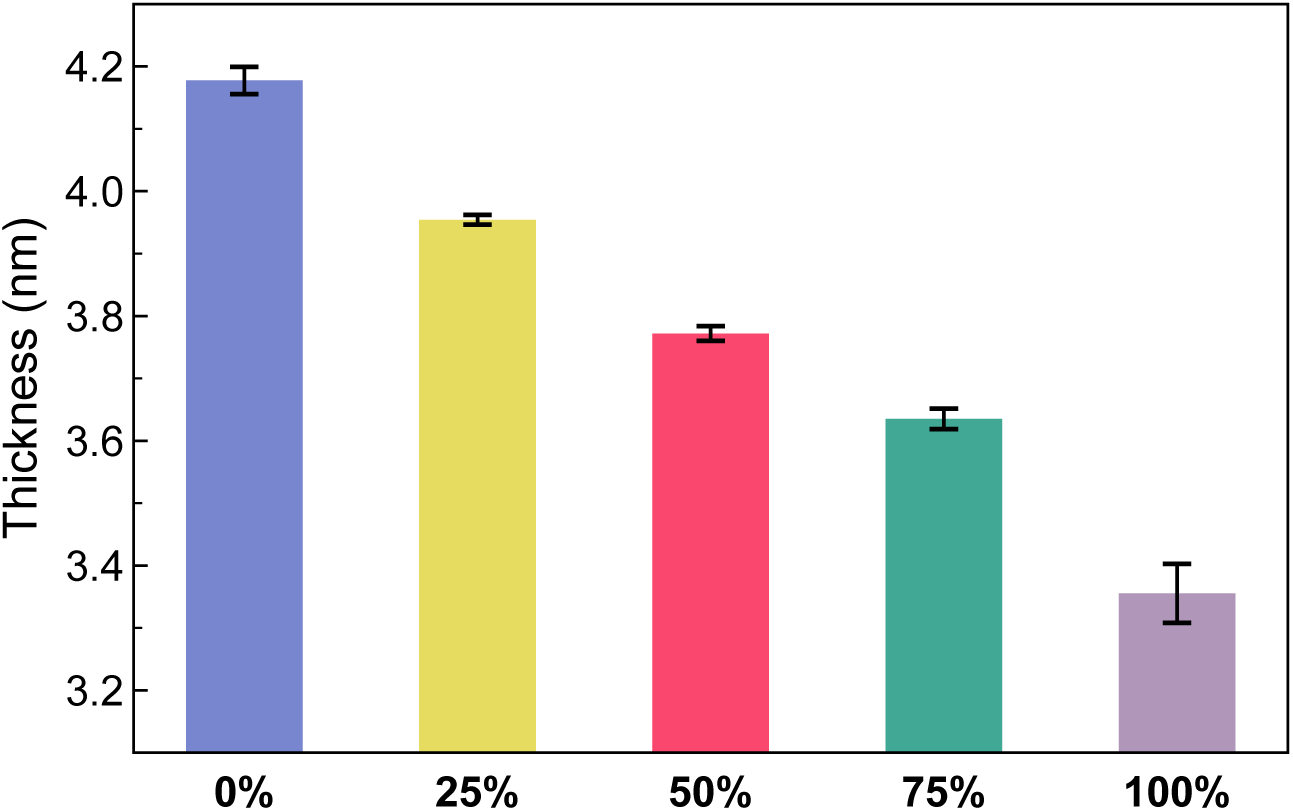
Membrane thickness as a function of oxidation level. Thickness values were calculated using `FATSLiM` and averaged over the final 50 ns of each trajectory, followed by averaging over three independent replicas. Error bars represent the standard deviation across replicas. Color coding: 0% (blue), 25% (yellow), 50% (red), 75% (green), 100% (lilac) PoxnoPC.

This behavior is consistent with the APL analysis, where the oxidized lipid species exhibit larger lateral area than their non-oxidized counterparts. Together, these results suggest that oxidation perturbs bilayer organization in a coupled manner, increasing lateral expansion while reducing the inter-leaflet thickness. Such changes are consistent with a progressive loosening of membrane packing and restructuring of the bilayer architecture upon oxidation.

### 3.3. Membrane Density and Bilayer Structural Rearrangement

To quantify the oxidation-induced redistribution of lipid mass along the bilayer normal, normalized number–density profiles were computed for the headgroup, glycerol-ester linkage, and acyl-tail regions of all membrane lipids using *ρ*_norm_(*z*) = *ρ*(*z*)*/ ρ*(*z*)*dz*. Figure 7a shows the interfacial headgroup density profiles. For the native membrane (0% PoxnoPC), two symmetric and well-defined peaks (≈ 65.5%) appear at *z* ≈ ±2.2 nm, consistent with a thick, well-packed interface. As the level of POPC oxidation increases, both the peak height and separation decrease. At 25 − 50% PoxnoPC, the peaks are moderately reduced to ≈ 56 − 52% of their maximum value. They shift slightly toward the center, indicating early thinning and softening of the membrane while retaining a relatively well-defined interface. At higher oxidation (75 − 100%), the peaks broaden and shift further inward (*z* ≈ ±1.7 nm) with intensities of ≈ 46%, reflecting a clear reduction in bilayer thickness and weakened inter-lipid packing due to the shortened, more flexible oxidized chains. A similar oxidation-dependent progression is observed for the glycerol–ester linkage region (Figure 7b). In 0 − 50% oxidation, the glycerol–ester peaks remain sharp and well-separated from the bilayer interior. At 75 − 100% PoxnoPC, these peaks broaden and move closer to the midplane, indicating partial penetration of oxidized fragments toward the core and disruption of the headgroup–tail boundary. Together with the headgroup results, this trend highlights the loss of interfacial definition and a corresponding increase in membrane fluidity under high peroxidation conditions. Such behavior has been reported in prior simulations of aldehyde-containing phospholipids, where truncated *sn*-2 chains exhibit enhanced flexibility and inward reorientation toward the bilayer core, accompanied by increased core hydration, as explicitly demonstrated in atomistic simulations of oxidized PC membranes [9, 54–56]. The hydrophobic-core density profiles (Figure 7c) further illustrate this restructuring. The native bilayer shows two maxima flanking a deep central minimum, characteristic of a thick, well-ordered hydrophobic core. With increasing oxidation, the density maxima increase in magnitude while shifting toward the midplane, and the central minimum rises, indicating that oxidized *sn*-2 chains reorient toward the bilayer center partially occupying the normally low-density midplane. As a results, the hydrophobic thickness is reduced and the membrane interior becomes more compressible.

**Figure 7:**
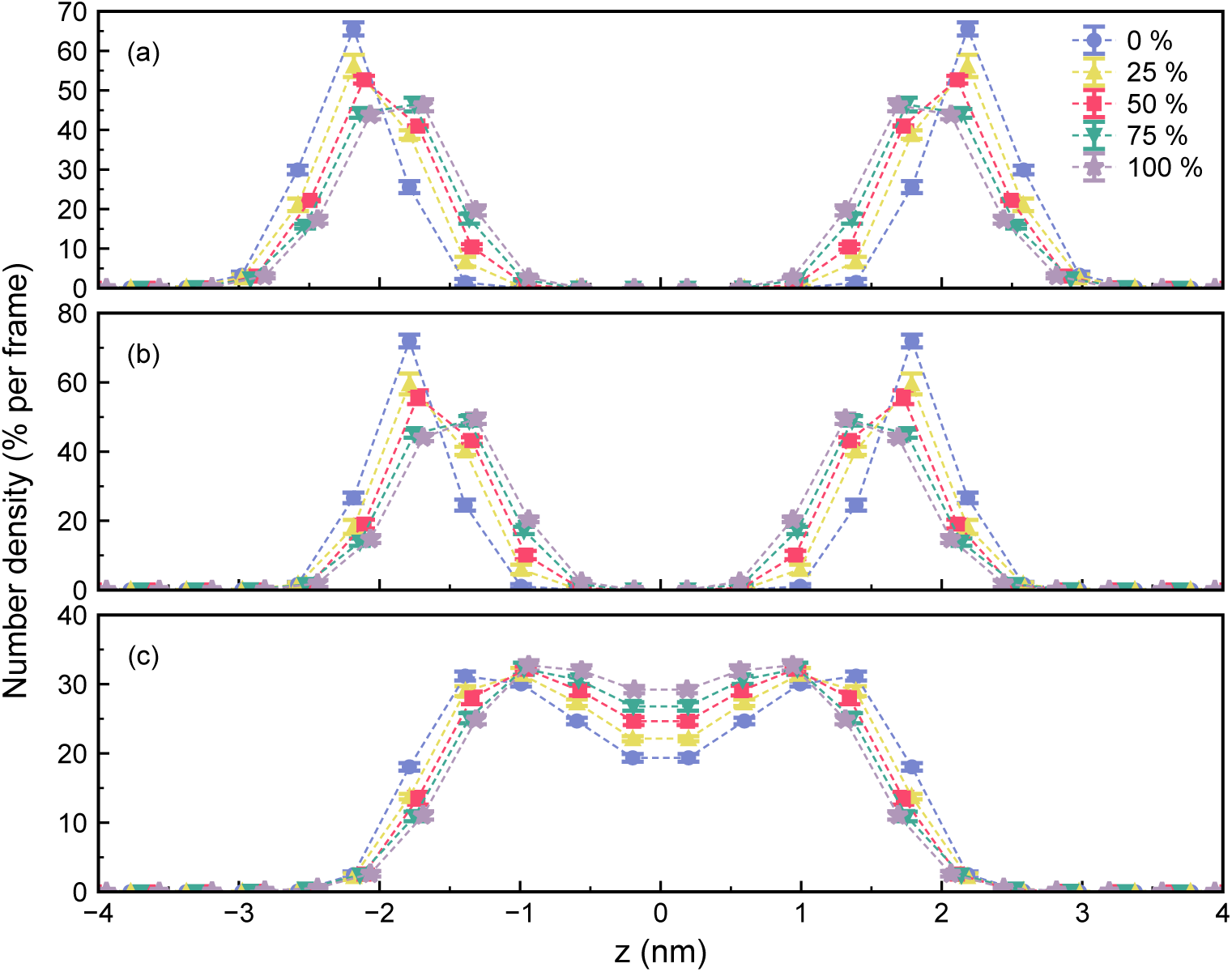
Normalized number-density profiles of membrane lipids at different POPC oxidation levels. (a) Headgroup region, (b) glycerol–ester linkage region, and (c) hydrophobic acyl-tail region. All profiles show symmetric distributions across the bilayer midplane (*z* = 0), and oxidation progressively shifts density inward, reflecting reduced bilayer thickness, weakened interfacial definition, and tighter but shorter hydrophobic core packing. Color coding: 0% (blue), 25% (yellow), 50% (red), 75% (green), 100% (lilac) PoxnoPC.

Similar increases in midplane density have been linked to enhanced hydration of the bilayer interior in membranes containing aldehyde-terminated oxidized lipids, where polar tail fragments promote water defects and permeability [7, 9, 54, 55].

Together, these effects indicate a more disordered and permeable membrane interior under high peroxidation.

### 3.4. Oxidation-Dependent Changes in Lateral Lipid Organization

Membrane composition also modulates how oxidation affects protein–bilayer interactions. Viral envelopes contain a heterogeneous mixture of phospholipids, glycolipids, and CHL. CHL is particularly important because it regulates lipid packing, membrane rigidity, and the formation of raft-like nanodomains [57–59], which can locally concentrate viral fusion and entry proteins, including the SARS-CoV-2 spike and its host receptors [58, 60]. Such lateral heterogeneity can buffer the bilayer against mild oxidative perturbations or, conversely, redistribute oxidized lipids between ordered and disordered domains, thereby modulating the local impact of peroxidation on membrane structure and permeability [9, 61]. Therefore, analyzing lipid clustering provides a more granular perspective on oxidation effects than can be obtained from bulk membrane properties alone.

To quantify how POPC oxidation reshapes lateral membrane organization, we evaluated clustersize distributions of membrane lipids at each oxidation level. Clusters were defined as groups of lipids whose reference atoms (P for phospholipids, O for CHL) lie within the RDF-derived cutoff distance (Section 2.3). Representative snapshots illustrating the identification of mixed POPC–PoxnoPC–CHL clusters, including the treatment of periodic boundary conditions, are shown in Figure 8. Cluster sizes were classified into three groups: **proto-clusters** (3 − 4 lipids), **small clusters** (5 − 10 lipids), and **large clusters** (*>* 10 lipids). These categories distinguish short-range contacts from cooperative nanoscale assemblies that resemble lipid-raft precursors. To isolate the direct consequences of acyl-chain oxidation, we first examined clustering exclusively within the phosphatidylcholine (POPC and its oxidized product, PoxnoPC) population. As shown in Table 3, increasing PoxnoPC content leads to a gradual reduction in both proto-clusters (3 − 4 lipids) and small clusters (5 − 10 lipids). Large clusters remain rare across all oxidation levels (< 0.5%), consistent with the absence of strong packing promoters in PC-only membranes. These results suggest that oxidized lipids disrupt lipid self-association, leading to smaller and more transient clusters.

**Figure 8:**
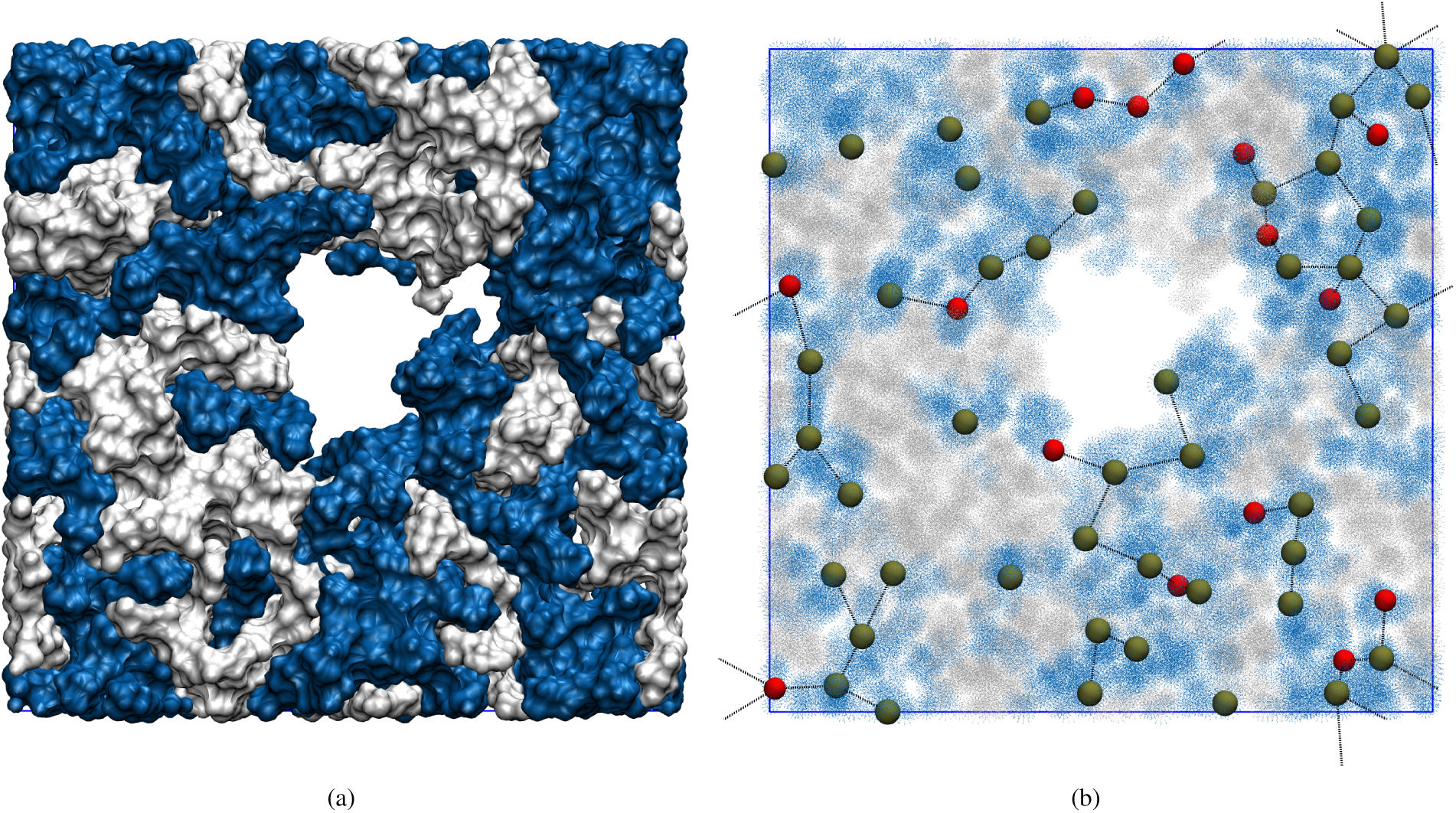
Representative visualization of lipid clustering in a CHL-containing oxidized membrane (POPC + PoxnoPC + CHL). (a) Top view of a single membrane leaflet shown in surface representation. Lipids belonging to a selected cluster are highlighted in blue, while the remaining membrane is shown in white. (b) Detailed atomistic representation of the same snapshot. Phosphorus atoms of POPC and PoxnoPC are shown as tan van der Waals spheres, and CHL oxygen atoms are shown as red van der Waals spheres. Lipids whose reference atoms lie within the clustering cutoff distance (*r*_cut_ = 0.9 nm) are connected by dashed lines and assigned to the same cluster. The simulation box is indicated by the blue rectangle. Periodic boundary conditions were applied during clustering analysis; therefore, nearest neighbors across periodic images are included, as illustrated by dashed connections between atoms in the primary box and their periodic images.

**Table 3:** Percentage distribution of POPC and PoxnoPC molecules participating in proto-clusters (3 − 4 lipids), small clusters (5 − 10 lipids), and large clusters (> 10 lipids) at different POPC oxidation levels. Values represent mean ± standard deviation from three independent replicas.

|  | **POPC+PoxnoPC** |  |  |
| --- | --- | --- | --- |
|  | proto cluster | small cluster | large cluster |
| 0% | 23.3 ± 0.7 | 8.4 ± 0.7 | 0.12 ± 0.05 |
| 25% | 22.5 ± 0.5 | 8.4 ± 0.7 | 0.06 ± 0.03 |
| 50% | 20.9 ± 0.7 | 9.0 ± 0.8 | 0.4 ± 0.1 |
| 75% | 17.9 ± 0.8 | 5.0 ± 0.4 | 0.02 ± 0.02 |
| 100% | 19.3 ± 0.7 | 6.3 ± 0.3 | 0.06 ± 0.03 |

**Table 4:** Percentage distribution of POPC, PoxnoPC, and CHL molecules participating in proto-clusters, small clusters, and large clusters at different POPC oxidation levels. Values represent mean ± standard deviation from three replicas.

|  | **POPC+PoxnoPC+CHL** |  |  |
| --- | --- | --- | --- |
|  | proto cluster | small cluster | large cluster |
| 0% | 14.2 ± 0.5 | 28.0 ± 0.8 | 39.7 ± 0.7 |
| 25% | 15.1 ± 1.5 | 27.9 ± 1.2 | 37.8 ± 1.3 |
| 50% | 15.1 ± 1.5 | 25.3 ± 0.9 | 35.4 ± 1.4 |
| 75% | 17.9 ± 0.9 | 30.7 ± 1.4 | 24.2 ± 1.1 |
| 100% | 15.9 ± 0.7 | 31.1 ± 1.0 | 25.9 ± 0.7 |

In the native membrane, CHL is a potent promoter of domain formation, yielding a dominant population of large clusters. At low-to-moderate oxidation (25 − 50%), these domains remain prevalent and changes are modest, suggesting that sterol–phospholipid interactions buffer the membrane against early-stage disorder. This buffering capacity is overwhelmed at higher oxidation (75 − 100%), however, where large cluster domains are substantially reduced and their components are redistributed into smaller aggregates, indicating that extensive peroxidation disrupts sterol-stabilized packing. Thus, CHL-rich domains preserve lateral order under mild oxidative stress, severe oxidation surpasses this effect, leading to the disruption of large clusters.

Overall, oxidation selectively disrupts large clusters even in CHL-rich membranes, and this structural disruption is correlated with the weakening of spike–membrane coupling at high oxidation levels.

## 4. Conclusions

Reactive oxygen species have dual effects on enveloped viruses: mild levels of lipid peroxidation can be tolerated, or even exploited, during infection, whereas severe oxidative damage disrupts the viral envelope and neutralizes the pathogen. In this work, we used atomistic MD simulations of ERGIC-like SARS-CoV-2 membranes to quantify how POPC oxidation modulates spike anchoring stability and surrounding bilayer properties. Umbrella sampling revealed a statistically significant dose-dependent effect on the spike detachment free energy. Oxidation of POPC up to 25 − 75% (corresponding to ≲ 42% oxidation of all PO-type phospholipids) led only to reductions in the TM+CT extraction barrier that were not statistically distinguishable from the non-oxidized reference within the sampling uncertainties of the present simulations. In contrast, the fully oxidized POPC condition, which corresponds to ≈ 55% of all glycerophospholipids, reduced the anchoring free energy by ≈ 23%, demonstrating that extensive peroxidation is sufficient to substantially weaken spike-membrane coupling even in the absence of direct chemical damage to the protein.

Membrane-structural analysis further revealed the mechanism of this destabilization. POPC peroxidation increases the area per lipid of the oxidized membrane components while progressively decreasing bilayer thickness, indicating lateral expansion and membrane thinning upon oxidation. POPC peroxidation also substantially decreases acyl-chain order in both *sn*-1 and *sn*-2 chains across all phospholipid species, with a stronger effect on *sn*-2 due to the truncated, kinked tail of PoxnoPC. Normalized density profiles show that oxidation thins the bilayer, broadens the headgroup and glycerol-ester distributions, and shifts both interfacial and tail densities toward the bilayer midplane, consistent with a softer, more compressible hydrophobic core. This loss of local order is also reflected at a larger scale, as oxidized POPC weakens cooperative packing. In PC-only membranes, this reduces the fraction of lipids in even small clusters, while in POPC+PoxnoPC+CHL mixtures it progressively diminishes large clusters at high oxidation levels. CHL thus delays, but does not prevent, the breakdown of ordered nanodomains under severe oxidative stress. Together, these changes, i.e., a thinner, more disordered, and less cohesive hydrophobic core, reduce hydrophobic matching and lower the energetic cost of extracting the TM helix and its CT domain from the bilayer.

Our findings connect atomistic membrane physics to the experimentally observed dose-dependent response of enveloped viruses to oxidative stress. Low levels of oxidation, comparable to those arising from intrinsic inflammatory ROS production, preserve spike anchoring and, therefore, are unlikely to strongly impair viral entry. In contrast, high oxidative doses – such as those from cold atmospheric plasma or ozone-based antiviral treatments – drive substantial disruption of membrane structure and lateral organization, providing a plausible molecular mechanism for the oxidative inactivation of SARS-CoV-2. This work highlights lipid peroxidation as a physically grounded contributing factor in antiviral mechanisms: extensive oxidation of POPC alone is sufficient to measurably weaken spike anchoring in a realistic multicomponent membrane. The results underscore the importance of membrane composition and lipid chemical damage as key regulators of coronavirus infectivity and motivate future studies that couple oxidized lipid fingerprints with spike conformational dynamics, fusion kinetics, and experimentally accessible markers of viral envelope integrity.

## Conflicts of interest

The authors declare no conflict of interest.

## Data availability

Supporting Information and the input files, parameters files and topology files as well as all figures and tables are available at: https://github.com/maryamghasemitarei/Impact-of-viral-membrane-oxidation-on-SARS-CoV-2-spike-protein-tail-stability

## Acknowledgments

We thank Annemie Bogaerts and Mohammad Reza Ejtehadi for their constructive feedback and expert guidance throughout the development of this work. T.A.N and M.G. acknowledge financial support from the NextGeneration EU Instrument of the European Union through the Academy of Finland (Grant No. 353298). Computational resources were provided by CSC IT Center for Science, Finland, and the RAMI RawMatters Finland Infrastructure. M.K. thanks Foundation PS, and the Research Council of Finland (Flagship of Advanced Mathematics for Sensing Imaging and Modelling grant 358944) for financial support.

## Notes

https://github.com/maryamghasemitarei/Impact-of-viral-membrane-oxidation-on-SARS-CoV-2-spike-protein-tail-stability/tree/main/project-files

### Competing Interest Statement

The authors have declared no competing interest.

