## Supplementary Information for "Impact of viral membrane oxidation on SARS-CoV-2 spike protein transmembrane anchoring stability"

### Structural Equilibration of Protein–Membrane Systems

*RMSD convergence of the spike TM–CT region*

To verify that all membrane systems were well equilibrated before property analysis and free-energy calculations, the structural stability of the spike protein embedded in each oxidation level was evaluated using the root-mean-square deviation (RMSD) of $C_\alpha$ of the transmembrane and cytoplasmic-tail region (TM–CT). RMSD values were computed on the 150 ns $NPT$ production simulations for all three replicas per oxidation level, after alignment to the initial configuration of the membrane-embedded protein segment. RMSD traces are shown in figures S1 for oxidation levels of 0%, 25%, 50%, 75% and 100%.

Across all systems, the RMSD exhibits an initial relaxation phase within the first $\approx 50$ ns, followed by a stable plateau, indicating the absence of large conformational drift or disruption of the secondary-structure. The final 50 ns of each trajectory (plateau region) were therefore selected to averaging all membrane structural properties (order parameters, number-density profiles, lipid clustering) to ensure analysis of fully equilibrated bilayer environments. Additionally, RMSD stability confirms that the oxidation-induced effects observed in free-energy and membrane-organization results originate from changes in the lipid structure rather than from significant rearrangements of the TM–CT region itself.

### Lipid-Specific Acyl-Chain Order Parameters

*sn-1 and sn-2 order profiles for each phospholipid class*

To complement the membrane-averaged order parameters reported in the main text, figure S2 shows the chain-resolved deuterium order parameters ($S_{\rm CD}$) separately for each phospholipid species present in the viral envelope: POPC/PoxnoPC, POPE, POPI, and POPS. Results are shown for both acyl chains (*sn*$-1$ and *sn*$-2$), and for all five oxidation levels (0, 25, 50, 75, and 100% POPC oxidation). For the *sn*$-2$ chain in the POPC/PoxnoPC plot, only the conserved C1–C7 segment

*Corresponding author
*Email addresses:* (Maryam Ghasemitarei), (Cristina Gyursánszky), (Mikko Karttunen), (Tapio Ala-Nissila )

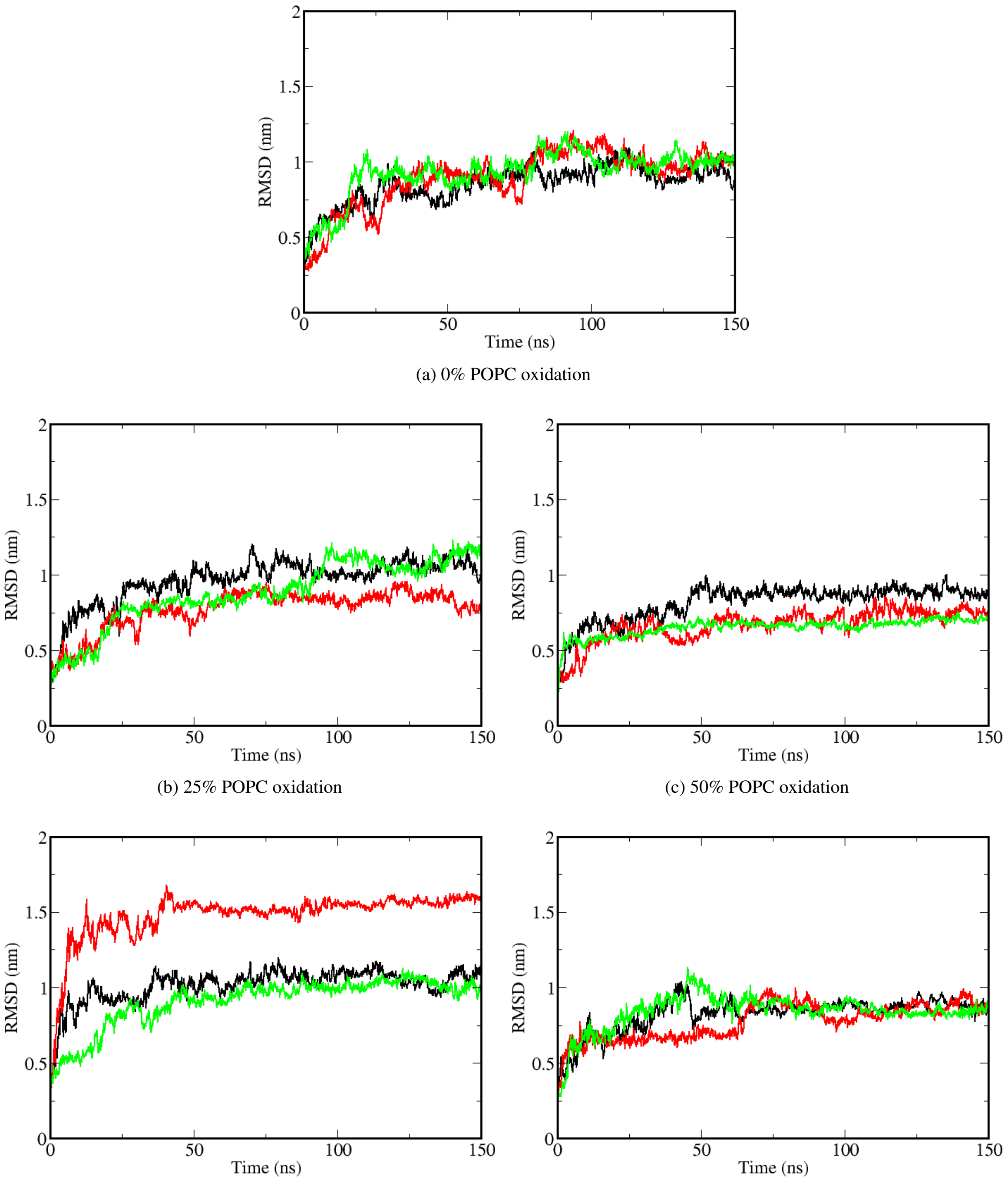


(a) 0% POPC oxidation

(b) 25% POPC oxidation

(c) 50% POPC oxidation

(d) 75% POPC oxidation

(e) 100% POPC oxidation

Figure S1. RMSD profiles of the spike TM–CT segment at different POPC oxidation levels. Each panel shows the RMSD over 150 ns for three replicas, confirming stable equilibration of the protein before production analyses.

is reported due to oxidation-induced truncation of PoxnoPC beyond C7. Across all lipid types, increasing PoxnoPC content results in a systematic decrease in $S_{\mathrm{CD}}$, demonstrating that oxidation-induced disorder propagates beyond the chemically modified lipids into their surrounding environment. This effect is most pronounced for $sn{-}2$ chains and for lipids proximal to oxidized POPC, reflecting loss of tight packing and increased free volume in the hydrophobic core.

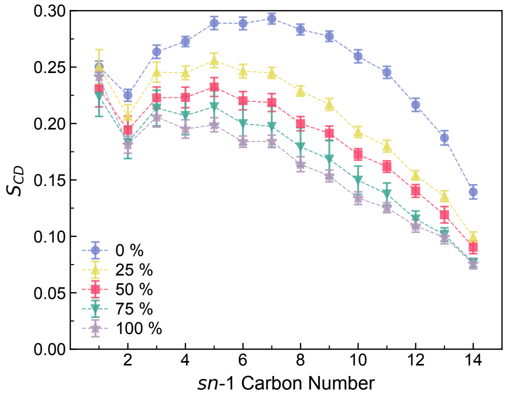


(a) POPC+PoxnoPC, *sn*-1

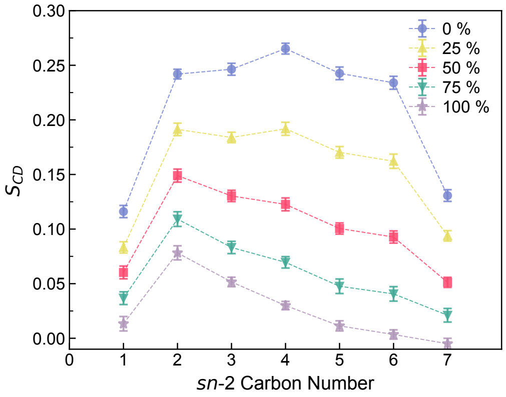


(b) POPC+PoxnoPC, *sn*-2

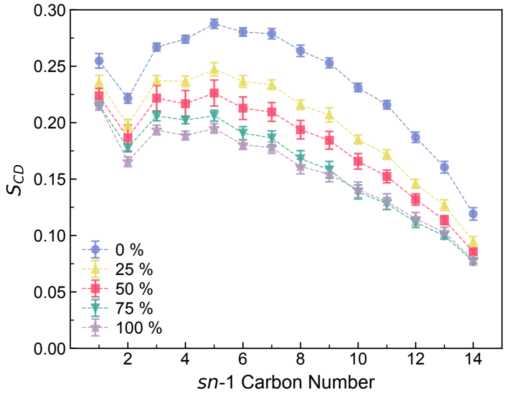


(c) POPE, *sn*-1

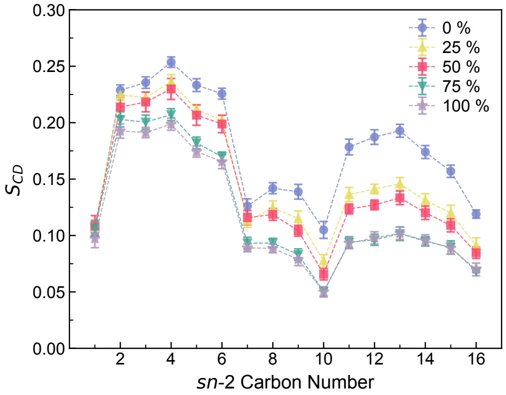


(d) POPE, *sn*-2

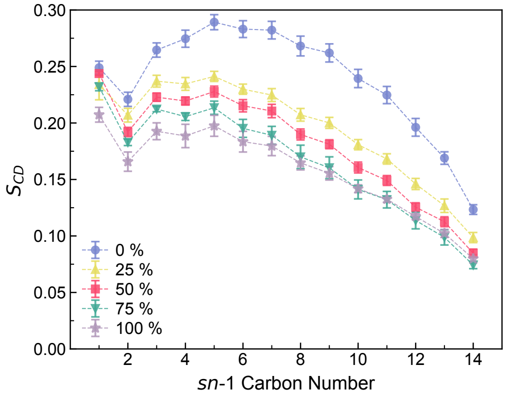


(e) POPI, *sn*-1

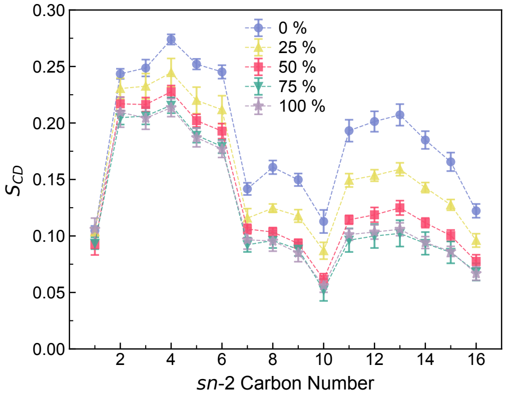


(f) POPI, *sn*-2

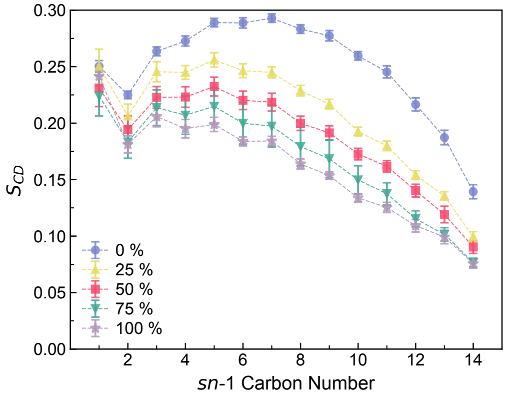


(g) POPS, *sn*-1

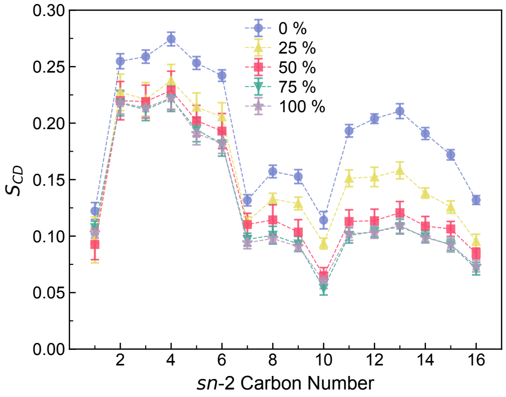


(h) POPS, *sn*-2

Figure S2. Chain-resolved $S_{\mathrm{CD}}$ profiles for the $sn-1$ and $sn-2$ acyl chains of each phospholipid type at different POPC oxidation levels (0% blue, 25% yellow, 50% red, 75% green, 100% lilac). Each row shows one lipid class.
